# The stress-induced protein NUPR1 orchestrates protein translation during ER-stress by interacting with eIF2α

**DOI:** 10.1101/2020.02.18.954115

**Authors:** Maria Teresa Borrello, Patricia Santofimia-Castaño, Marco Bocchio, Angela Listi, Nicolas Fraunhoffer, Philippe Soubeyran, Eric Chevet, Christopher Pin, Juan Iovanna

**Author notes:** Corresponding author address: Juan Iovanna.; Borrello Maria Teresa, Centre de Recherche en Cancérologie de Marseille (CRCM), INSERM U1068, CNRS UMR 7258, Aix-Marseille Université and Institut Paoli-Calmettes, Parc Scientifique et Technologique de Luminy, 163 Avenue de Luminy, 13288 Marseille, France.; Mobil.

## Abstract

NUPR1 is a stress response protein overexpressed upon cell injury in virtually all organs including the exocrine pancreas. Despite NUPR1’s well established role in the response to cell stress, the molecular and structural machineries triggered by NUPR1 activation remain largely unknown. In this study, we uncover an important role for NUPR1 in participating in the unfolded protein response pathway and the endoplasmic reticulum stress response. Biochemical results, confirmed by ultrastructural morphological observation, revealed alterations in the UPR in acinar cells of germline-deleted NUPR1 murine models, consistent with the inability to restore general protein translation. Bioinformatical analysis of NUPR1 interacting partners showed significant enrichment in translation initiation factors, including eukaryotic initiation factor (eIF) 2α. Co-immunoprecipitation and proximity ligation assays both confirmed interaction between NUPR1 and eIF2α and its phosphorylated form (p-eIF2α). Our. Moreover, our data also suggest loss of NUPR1 in cells results in maintained eIF2α phosphorylation and evaluation of nascent proteins by (peIF2α), and click chemistry revealed that NUPR1-depleted PANC-1 cells displayed a slower post stress protein translational recovery compared to wild-type. Combined, this data proposes a novel role for NUPR1 in the integrated stress response pathway, at least partially through promoting efficient PERK-branch activity and resolution through a unique interaction with eIF2α.

**Significance:** In the pancreas, NUPR1 is required for a resolution of the ER stress response. During ER stress response, NUPR1 binds both eIF2α allowing for its dephosphorylation and restoration of new protein synthesis.

**Highlights:** Biochemical analysis revealed a general reduction in the protein expression of downstream mediators of the unfolded protein response in the pancreas of mice lacking *Nupr1*. This finding suggests a novel role for NUPR1 in the UPR/ER stress response.

Ultrastructural analysis of pancreata revealed reduced morphological alterations in tunicamycin-treated *Nupr1*^-/-^ mice compared to *Nupr1*^+/+^ mice consistent with a maintained block in general protein translation.

Co-immunoprecipitation of tagged NUPR1 confirmed a novel interaction with eIF2α. Depletion of NUPR1 prolonged phosphorylation of eIF2α, suggesting it may be involved in attenuation of the PERK branch of the UPR.

NUPR1-depleted PANC-1 cells displayed a slower recovery of protein translation following UPR activation

## Introduction

NUPR1, also known as p8 or Com1, is an intrinsically-disordered protein first identified during the onset of pancreatitis [1]. We and others demonstrated NUPR1 is transiently induced in almost all organs and cells in response to a variety of injuries [2–6] including minimal stresses such as the renewal of culture medium [7]. While it is clear NUPR1 acts as an essential element during the stress cell response, protecting cells from genotoxic or oxidative injury [8–11], the mechanisms by which NUPR1 acts still need to be elucidated. The highest expression of NUPR1 has been reported in pancreatic acinar cells following induction of pancreatitis. Acinar cells are highly enriched for endoplasmic reticulum (ER) due to their having the highest rate of protein synthesis among all cell types [12,13]. A major function of the ER involves folding and post-translational modification of secreted and integral membrane proteins. Also, the ER maintains homeostasis between folded and unfolded proteins [14,15] and disturbance of these physiological ER activities leads to a cell stress response implicated in a variety of pathological states [14,16,17]. Several pieces of evidence indicate NUPR1 is involved in the onset of ER stress but its role in this context remains largely unexplored [18,19].

ER stress can be activated by a number of events including accumulation of unfolded proteins in the ER, subsequently triggering several signaling pathways that, together, are termed the unfolded protein response (UPR). The ultimate goal of the UPR is to resolve homeostatic imbalance between folded and unfolded proteins. If protein homeostasis is not restored, the UPR triggers apoptosis to safely dispose of damaged cells. The UPR is comprised of three main branches - PKR-like ER kinase (PERK), inositol-required enzyme 1 (IRE1) and activation transcription factor 6 (ATF6). In the absence of stress, these molecules are bound to the chaperone BiP (GRP78 or HSPA5). When excessive protein load occurs, BiP dissociates from PERK, IRE1 and ATF6 leading to their activation [20]. Activation of PERK leads to phosphorylation of the eukaryotic initiation factor 2α (eIF2α), which is a critical regulator of protein translation. Phosphorylated eIF2α prompts a dramatic reduction in protein translation to limit cellular amino-acid consumption. However, several mRNA transcripts elude the translation block, including Activating Transcription Factor 4 (ATF4). ATF4 increases cell survival by promoting the expression of genes involved in protein folding, amino-acid import and biosynthesis of aminoacyl-transfer RNAs [21]. When protein homeostasis is restored, GADD34/PPP1R15A, which is activated by ATF4, promotes eIF2α dephosphorylation, thereby restoring normal protein synthesis [22]. Prolonged or excessive ER stress activation leads to programmed cell death, mainly mediated by the activation of the CCAAT/enhancer binding protein (C/EBP) homologous protein (CHOP) [16,22], which promotes apoptosis via the intrinsic mitochondrial pathway.

The goals of this study were to determine if the absence of NUPR1 altered the ER stress response and UPR in the pancreas and determine a possible mechanism by which NUPR1 affected this pathway. We report that NUPR1 constitutes an important element of the UPR activation. Our data shows in the absence of NUPR1 in the pancreas, phosphorylation of eIF2α is maintained, thereby preventing restoration of protein translation and, likely, contributing to increased tissue damage. We provide evidence that NUPR1’s directly interacts with eIF2*α*, and may be required for its dephosphorylation and restoration of proper protein translation. This is a completely novel function for NUPR1 and, for the first time, links these two critical factors involved in the integrated stress response.

## Results

### Biochemical evaluation of ER-stress activators in pancreatic acinar cells of *Nupr1*^*+/+*^ and *Nupr1*^*-/-*^ mice

Recently, we reported mice lacking *Nupr1* (*Nupr1*^*-/-*^) showed a significant downregulation in genes related to ER stress in the pancreas [18] suggesting NUPR1 is necessary for mediating a correct UPR activation. To test the importance of NUPR1 in the onset of the ER-stress response, pharmacological activation of stress was initiated by a single injection of tunicamycin (Tun, 1.0 µg/g) in *Nupr1*^*-/-*^ and *Nupr1*^*+/+*^ mice. Tun induces ER stress by inhibiting protein *N*-Glycosylation, thereby preventing correct protein folding [23]. Sixteen hours after Tun administration, whole pancreatic protein extracts were collected and the expression levels of the major UPR mediators of the PERK and IRE1-α branches evaluated by western blot and qPCR (Figure 1). PERK accumulation and phosphorylation appeared similar between the two genotypes. Consistent with activation of PERK signaling, increased expression of ATF4 was observed in both *Nupr1*^*-/-*^ and *Nupr1*^*+/+*^ following activation of cell stress (Figure 1). However, the increase in ATF4 expression was reduced in *Nupr1*^*-/-*^ mice at both protein and mRNA levels (Figure 1G) compared to *Nupr1*^*+/+*^ tissue following Tun injection (*p <* 0.0001). Reduced ATF4 function was supported by the absence of CHOP protein in *Nupr1*^*-/-*^ pancreata (Figure 1A, B). Interestingly, while we observed reduced induction of *Chop* mRNA expression in *Nupr1*^*-/-*^ mice compared to *Nupr1*^*+/+*^ after stress (effect of Tun: *p<* 0.0001;; effect of genotype *p <* 0.0001;; interaction *p <* 0.0001;; *Nupr1*^*+/+*^ vs *Nupr1*^*-/-*^ in Tun condition: *p <* 0.0001, *post hoc* Sidak test, n=6), an increase in *Chop* mRNA was still observed (Figure 1G), suggesting protein translation may be affected. Similar analysis of another ATF4 target, *Gadd34*, indicated lower expression in *Nupr1*^*-/-*^ pancreata compared to *Nupr1*^*+/+*^ after stress induction. Since GADD34 is involved in a negative feedback loop to reduce PERK signaling, these findings suggest a possible delay in mechanisms involved of the resolution of ER-stress in *Nupr1*^*-/-*^.

**Figure 1.**
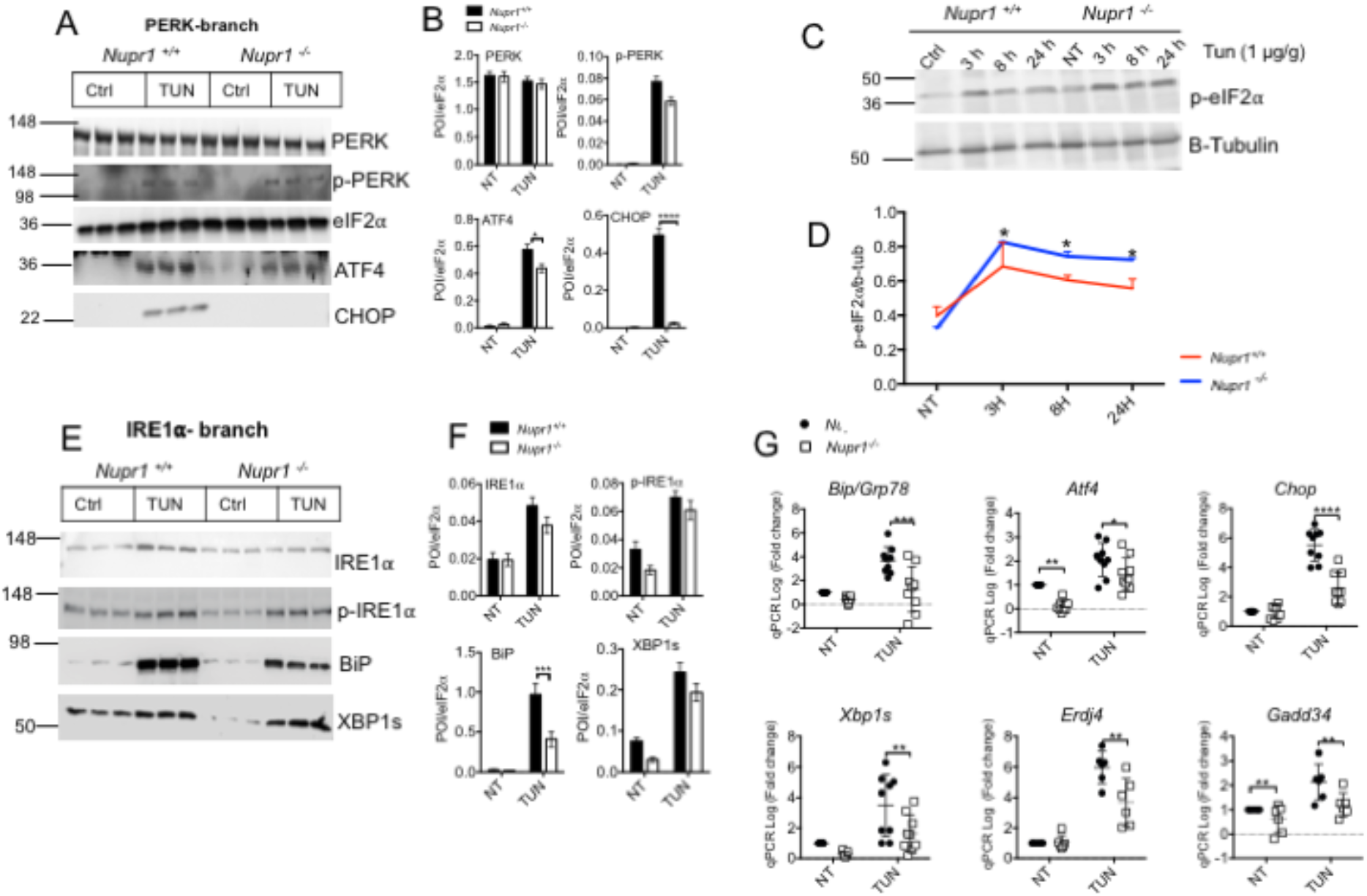
Biochemical evaluation of ER-stress related proteins after tunicamycin treatment in the pancreas. **(A)** *Nupr1*^+/+^ and *Nupr1*^*-*/-^ mice were injected with 1 µg of Tunicamycin or vehicle and 16 h (150 mM of D-glucose) post injection pancreas was extracted and tissue lysed. Image show representative western blots of the PERK branch activation and its quantification **(B)** using ImageJ software. Mean band intensity plotted ± SEM;; Significant differences were calculated by two-way ANOVA with *post hoc* Sidak test (n=3). (C) Representative western blotting of p-eIF2α of pancreatic lysates from *Nupr1*^+/+^ and *Nupr1*^*-*/-^ after Tun injections over a period of time (Ctrl, 3 h, 8 h, 24 h);; (D) quantification of (C) using ImageJ software. Mean band intensity plotted ± SEM;; Significant differences were calculated by two-way ANOVA with post hoc Sidak test (n=3). **(E)** Representative western blots of the IRE1*α* branch of the UPR of tissue lysates dissected from from *Nupr1*^+/+^ and *Nupr1*^*-*/-^ and quantification **(F)** using ImageJ software. Mean band intensity plotted ± SEM;; Significant differences were calculated by two-way ANOVA with *post hoc* Sidak test. Statistically significant differences between *Nupr1*^+/+^ and *Nupr1*^*-*/-^ mice are shown (*\*p* < 0.02, *\*\*p* < 0.01, \*\*\**p <* 0.001, *\*\*\*\*p<*0.0001). (**G**) RT-qPCR results of mRNA expression of BiP, *Atf6, Xbp1d, Atf4* and *Chop, Gadd34 and Erdj4* RNA was extracted from *Nupr1*^*+/+*^ and *Nupr1*^*-/-*^ mice treated with Tun or Vehicle for 16h. Significant differences were calculated by two-way ANOVA with *post hoc* Sidak test (n=6). Statistically significant differences are shown (*\*p* < 0.03, *\*\*p* < 0.001, *\*\*\*p* = 0.001, *\*\*\*\*p <* 0.0001).

The phosphorylation of eIF2α is a critical step in the cellular stress response, mediating a protein synthesis shut-off and GADD34 is directly involved in its dephosphorylation. Given reduced GADD34 expression, we next measured the levels of p-eIF2α up to 24 hours following Tun injection (Figure 1C,D). Three hours following Tun treatment similar levels of p-eIF2α are observed in *Nupr1*^*+/+*^ and *Nupr1*^*-/-*^ mice. However, at both 6 and 24 hours into Tun treatment, the level of p-eIF2α is significantly higher in *Nupr1*^*-/-*^ tissue, revealing a possible delay in the mechanism of ER-stress termination in *Nupr1*^*-/-*^.

Biochemical examination of IRE1*α* branch activation showed post-stress accumulation of IRE1*α* and phospho (p) IRE1*α* with no significant variation in both genotypes (Figure 1E-F). However, western blot analysis showed pancreatic expression of the chaperone BiP was almost halved in *Nupr1*^*-/-*^ mice compared to *Nupr1*^*+/+*^ mice (n=3). Expression of the *BiP* mRNA showed a consistent downregulation in *Nupr1*^*-/-*^ mice compared to *Nupr1*^*+/+*^ littermates (n=6). For both protein and mRNA, two-way ANOVA analysis revealed a significant effect of Tun (*p* < 0.0001 for protein expression and *p* < 0.001 for mRNA). Subsequent comparisons uncovered a significant difference in BiP between *Nupr1*^*+/+*^ and *Nupr1*^*-/-*^ mice after Tun administration (*p* < 0.0001 for protein and *p* = 0.0228 for mRNA, *post hoc* Sidak test) but not in control conditions.

BiP is a stress sensor of the UPR and an integral part of the ER quality control system. Upon UPR initiation, the translational efficiency of BiP is normally increased by 2-3 fold and it is regulated by several overlapping mechanisms [24]. Therefore, downregulation of BiP protein and mRNA expression could be correlated with a faulty UPR machinery in *Nupr1*^*-/-*^ mice. Similar levels of total IRE1α, p-IRE1α and spliced XBP1 (XBP1s) protein were observed in both *Nupr1*^*+/+*^ and *Nupr1*^*-/-*^ treated mice suggesting an effective activation of the IRE1α branch occurred. However, a significant difference was detected between the two genotypes when examining *Xbp1s* mRNA levels (Figure 1G). The increase in *Xbp1s* mRNA induced by stress in *Nupr1*^*-/-*^ mice was significantly lower compared to *Nupr1*^*+/+*^ mice (effect of Tun: *p <* 0.0001;; effect of genotype *p* = 0.003, with no interaction between the two factors;; *Nupr1*^*+/+*^ vs *Nupr1*^*-/-*^ in Tun condition: *p* < 0.0001 *post hoc* Sidak test, n=6). To validate the decrease of *Xbp1s* in *Nupr1*^*-/-*^ models, we evaluated *Erdj4* mRNA expression, *a* known target of XBP1s transcriptional regulation [25]. Consistent with reduced XBP1s function, *Erdj4* expression was decreased in *Nupr1*^*-/-*^ mice compared to the *Nupr1*^*+/+*^ littermates (*p =* 0.001). Taken together, these results indicate the absence of NUPR1 leads to a compromised activation of the UPR.

### NUPR1-deficiency could prevent initial ultrastructural alterations in murine pancreatic acinar cells after Tunicamycin induced cell stress

To determine the effects of deleting *Nupr1* on acinar cell morphology, we examined ultrastructural modifications of the ER in *Nupr1*^*+/+*^ and *Nupr1*^*-/-*^ mice 16 h or 36 h post-Tun treatment using transmission electron microscopy. Representative micrographs indicate non-treated exocrine pancreatic cells (Figures 2A and 2B) present with a normal ER, structurally ordered into thin, densely packed *cisternae* covered with ribosomes in both genotypes (arrows in insets of Figure 2A and 2B). Overall, cell ultrastructure was normal in *Nupr1*^*-/-*^ mice. Acinar cells showed regular polarized organization with visible mononucleated cells (Nu) and electron dense zymogen granules (zy). Sixteen hours post-Tun injection, acinar cells in *Nupr1*^*+/+*^ mice displayed dilated and expanded ER (Figure 2C and inset) with an almost complete loss of associated ribosomes in the perinuclear area and fewer electron dense zymogen granules. Expanded ER *cisternae* are a cellular hallmark of the UPR as ER membranes expand to alleviate the stress due to an excessive load of misfolded proteins. Increasing ER volume decreases the relative concentration of unfolded protein intermediates, increases the time for protein folding, and avoids aggregate formation [26]. While a decrease in zymogen granules was detected in *Nupr1*^*-/-*^ pancreata (Figure 2D), even 36 h after Tun treatment, little to no expansion of the ER was observed. As time progressed, damage to the ER was more aggravated in *Nupr1*^*+/+*^ mice, becoming fragmented (Figure 2E). Such ER abnormalities were observed in *Nupr1*^*+/+*^ samples in more than 40% of the analyzed acinar cells (12/30 cells from ten randomly selected fields of acquisition). Conversely, such ER dilation was almost completely absent in *Nupr1*^*-/-*^ samples (Figure 2F).

**Figure 2.**
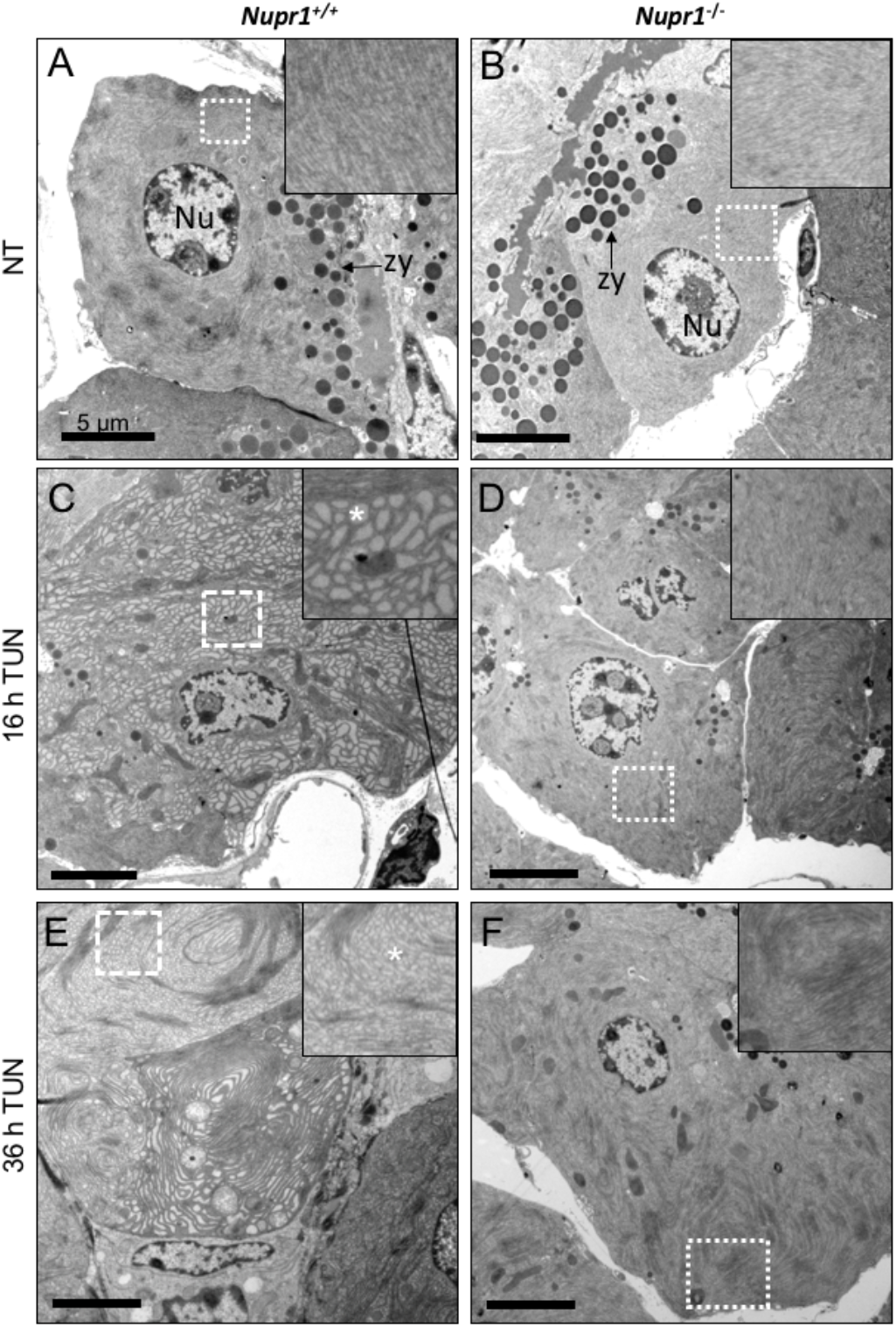
Electron micrographs of *Nupr1*^*+/+*^ and *Nupr1*^*-/-*^ murine pancreata after ER stress induction with tunicamycin (16 and 36 h). Representative transmission electron micrograph images of *Nupr1*^*+/+*^ and *Nupr1*^*-/-*^ pancreatic tissue of perfusion fixed mice injected intra peritoneally (IP) with vehicle (NT, A and B and insets) or Tunicamycin (TUN, 1.0 µg/g), at 16 (**C** and **D**) or 36 hours (**E** and **F**) post injection. Arrows point to the ER and asterisks indicate dilated ER cisternae. **Nu**=nucleus;; **zy** = zymogen granules.

The acinar cell phenotype observed in *Nupr1*^*-/-*^ mice suggests that translation of proteins may be generally affected in response to stress in these mice. Since an absence of NUPR1 is correlated to reduced *Gadd34* expression and limited expansion of the ER in *Nupr1*^*-/-*^ mice, we speculated NUPR1 is required for restoration of protein synthesis. To test this hypothesis, acinar cells were isolated and assessed in culture (Figure 3A). This process activates the UPR [27]. Morphological analysis of isolated acinar cells showed no obvious difference between *Nupr1*^*+/+*^ and *Nupr1*^*-/-*^ acini (Figure 3B). Quantification of amylase levels, however, showed a significant decrease in amylase protein in *Nupr1*^*-/-*^ cells (Figure 3C;; *p <* 0.05, *post hoc* Sidak test), suggesting protein translation was reduced in these animals. It is possible however, that lower amylase levels could reflect increased exocytosis in the absence of NUPR1. To examine this possibility, we measured secretion of amylase in *ex-vivo* pancreatic acini from *Nupr1*^*+/+*^ and *Nupr1*^*-/-*^ mice (Figure 3D). Following 30 min incubation with increasing concentrations of cerulein (analogue of cholecystokinin [28]) media amylase levels revealed a dose-dependent response for both genotypes (Figure 3D, effect of caerulein: *p < 0*.*012*, two-way ANOVA). Noteworthy, media-amylase levels were significantly lower in *Nupr1*^*-/-*^ acinar cultures compared to *Nupr1*^*+/+*^ cultures (Figure 3D, *p* < 0.0001, two-way ANOVA) supporting a model in which decreased amylase levels were due to either reduced translation or increased degradation in *Nupr1*^*-/-*^ acini.

**Figure 3.**
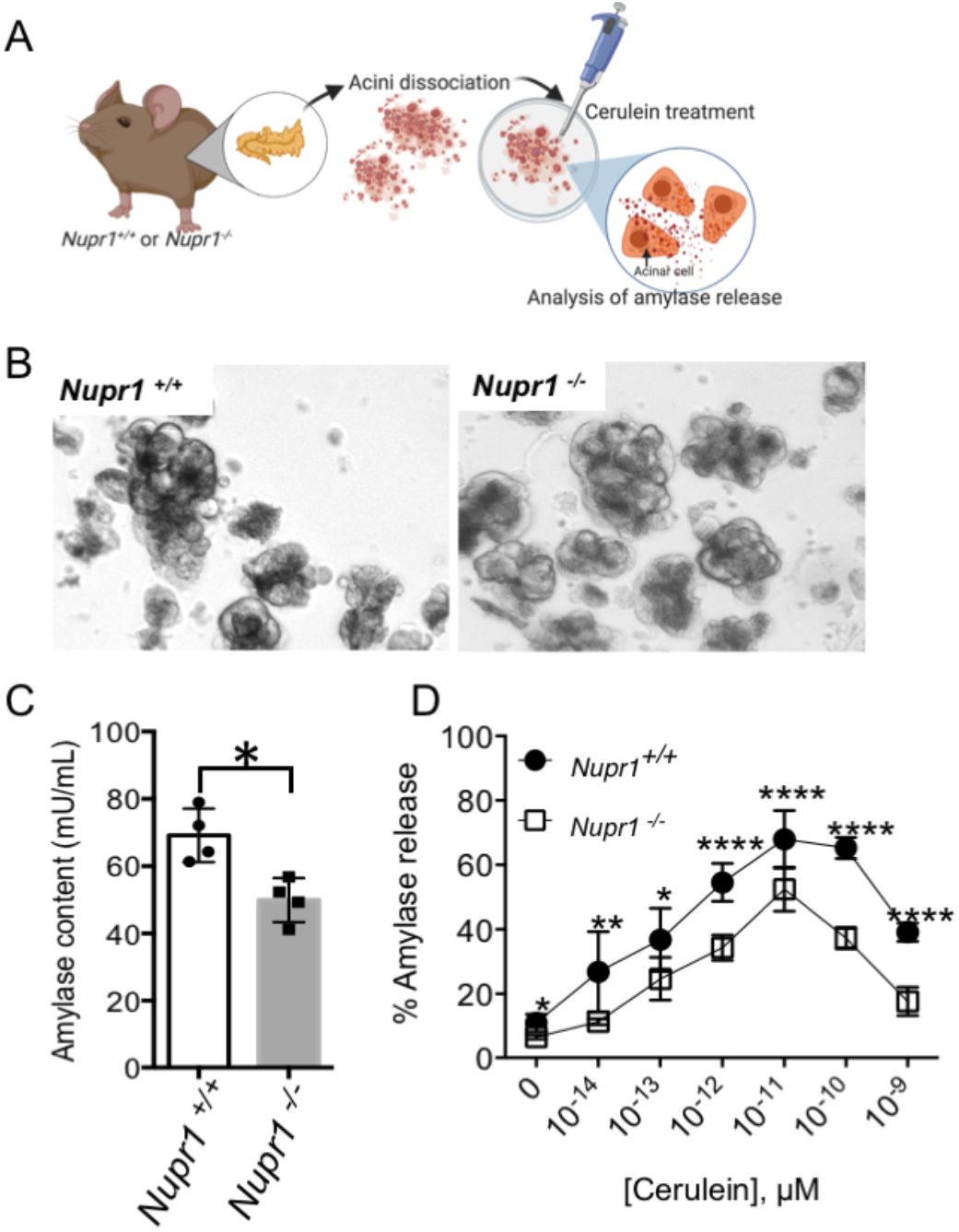
Quantification of amylase content and amylase release levels in acinar cells isolated from *Nupr1*^*+/+*^ and *Nupr1*^*-/-*^ murine pancreata. **(A)** Acini from *Nupr1*^*+/+*^ or *Nupr1*^*-/-*^ were dissociated and treated with increasing concentration of Cerulein. Amylase release onto the media was then measured. **(B)** Image of isolated acini from *Nupr1*^*+/+*^ or *Nupr1*^*-/-*^ mice. **(C)** Quantification of total amylse content in *Nupr1*^*+/+*^ and *Nupr1*^*-/-*^ expressed in mU/mL. Total amylase in *Nupr1*^*-/-*^ acini was significantly lower than from acini of *Nupr1*. Statistical significance was calculated with unpaired two-tailed *t*-test *\*p<* 0.04, n = 4). **(D)** Amylase release from isolated *Nupr1*^*+/+*^ or *Nupr1*^*-/-*^ acinar cells following stimulation with increased amounts of cerulein for 30 min. *Nupr1*^*+/+*^ derived acinar cells released higher percentage of Amylase compared to *Nupr1*^*-/-*^. Statistical significance was measured with two-way ANOVA with post hoc Sidak test;; \**p<*0.01, *\*\*p<*0.014, ****p< 0.0001;; n = 4 mice).

### Identifying the role of NUPR1 during ER stress response

Based on these observation, the loss of NUPR1 appears to affect restoration of protein translation in response to ER stress. To identify the molecular mechanisms through which NUPR1 may regulate the UPR, we performed immunoprecipitation for NUPR1 to determine putative interacting proteins. Flag-tagged NUPR1 was expressed in MiaPaCa-2 cells (Supplementary Figure 1) and immunoprecipitated 24 hours later under either normal conditions or following induction of ER-stress by glucose starvation or addition of 1 µM thapsigargin (TPS;; Figure 4). NUPR1-associated proteins were next identified by mass spectrometry (MS) resulting in 656 putative NUPR1-interacting proteins under normal conditions (Supplementary Table 1), and 1152 or 828 interacting proteins under glucose-starvation (Supplementary Table 2) and TPS conditions (Supplementary Table 3), respectively (Figure 4A). 577 proteins were common to all conditions, with 7 (non-treated), 365 (glucose-starvation) and 77 proteins (TPS) specific to the various conditions (Figure 4A). Bioinformatic analysis using the String protein-protein interaction database showed significant enrichment in translation initiation factors, supporting a model in which NUPR1 affects protein translation. Twenty-two out of a total of 142 proteins directly involved in translation initiation were identified under unstressed (non-treated) conditions (*p* = 4.14e-07;; Table 1). As expected for a role in protein translation during cellular stress, the number of NUPR1-interacting proteins increased to 73 (*p =* 5.19e-36) in glucose-starvation and 45 (*p=*4.76e-21) in TPS-treated cells (Table 1). This suggest an expanded role for NUPR1 in translational regulation during ER stress.

**Figure 4.**
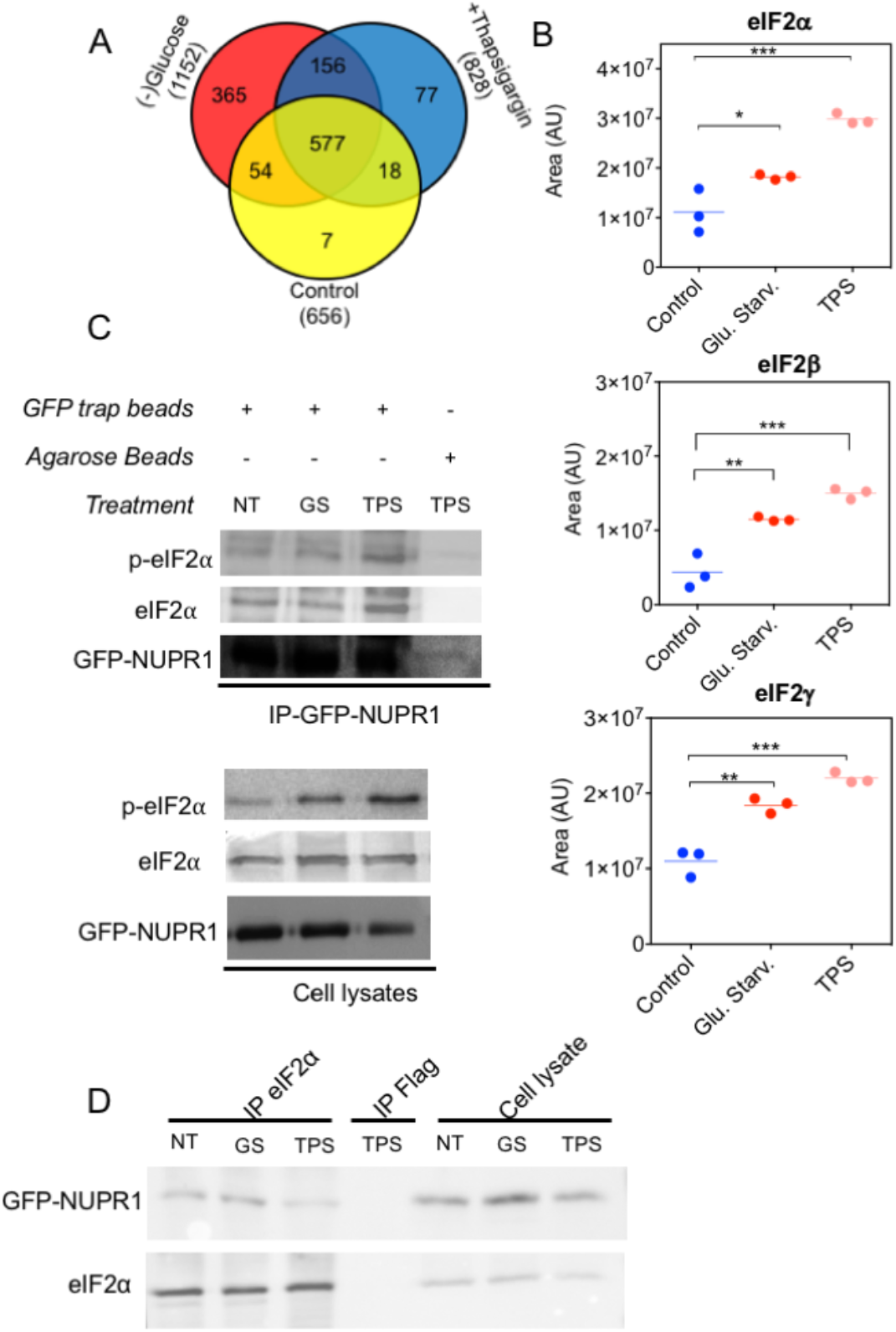
Identification of putative NUPR1 interacting proteins. **(A)** Venn diagram showing putative NUPR1-interacting proteins following glucose starvation or thapsigargin (TPS, 1 µM, 24 hours) treatment. **(B)** Quantification of mass spectrometry peak areas of NUPR1-associated eukaryotic initiation factors (n=3, one-way ANOVA, *\*p* = 0.02, *\*\*p =* 0.002, *\*\*\*p =* 0.0002). **(C)** Extracts from MiaPaCa-2 cells transiently transfected with GFP-NUPR1 were subjected to co-immunoprecipitation with GFP-trap^®^ beads or agarose beads followed by western blotting with the antibodies against p-eIF2α and eIF2α.Co-Immunoprecipitation assay of GFP-tagged-NUPR1 confirm p-eIF2α and eIF2α interaction. **(D)** Co-Immunoprecipitation of GFP-tagged-NUPR1 with eIF2α demonstrate the reciprocal interaction is effective.

The NUPR1 interactome included many translation initiation factors, which increased upon ER stress induction and included eIF2α (eIF2S1), eIF2*β* (eIF2S2), and eIF2*γ* (eIF2S3) (Figure 4B). These findings suggest a novel function for NUPR1 and provide a unique link between two important stress response proteins. Since our initial findings imply the levels of p-eIF2α are increased in Tun-induced *Nupr1*^*-/-*^ acinar tissue, we decided to confirm this interaction. First, co-immunoprecipitation was performed following expression of a GFP-tagged NUPR1 in MiaPaCa-2 cells (Figure 4C). The interaction between NUPR1 and eIF2α was confirmed in non-treated cells showing that NUPR1 will bind eIF2α even under non-stressed conditions. Upon glucose starvation and TPS treatment, this interaction was maintained and even increased in the presence of TPS, confirming mass spectrometry data. Next, we performed the reverse co-IP, pulling down GFP-tagged NUPR1 using eIF2*α* antibody (Figure 4D). To show this interaction took place within the cell, the association between NUPR1 and p-eIF2α/eIF2α, examined using a proximity ligation assay (PLA, Duolink^®^), which identifies molecular complexes that occur at distances <16 Å (Figure 5). Confocal fluorescent microscopy analysis revealed PLA positive foci under all conditions, suggesting NUPR1 interacts with eIF2α independent of cell stress. Importantly, this interaction is extranuclear, which would be expected for a direct interaction with eIF2α and p-eIF2α, and is the first direct evidence of an extra-nuclear function for NUPR1. PLA also revealed increased interaction following TPS treatment or glucose starvation (GS), with NUPR1-eIF2α and NUPR1-p-eIF2α interactions remaining in the perinuclear area (Figure 5A and 5C). These observations confirm NUPR1 interacts with both eIF2α and p-eIF2α and ER stress increases or stabilizes the interaction. Combined, our data suggests a novel role for NUPR1 regulating the translational machinery during ER stress.

**Figure 5.**
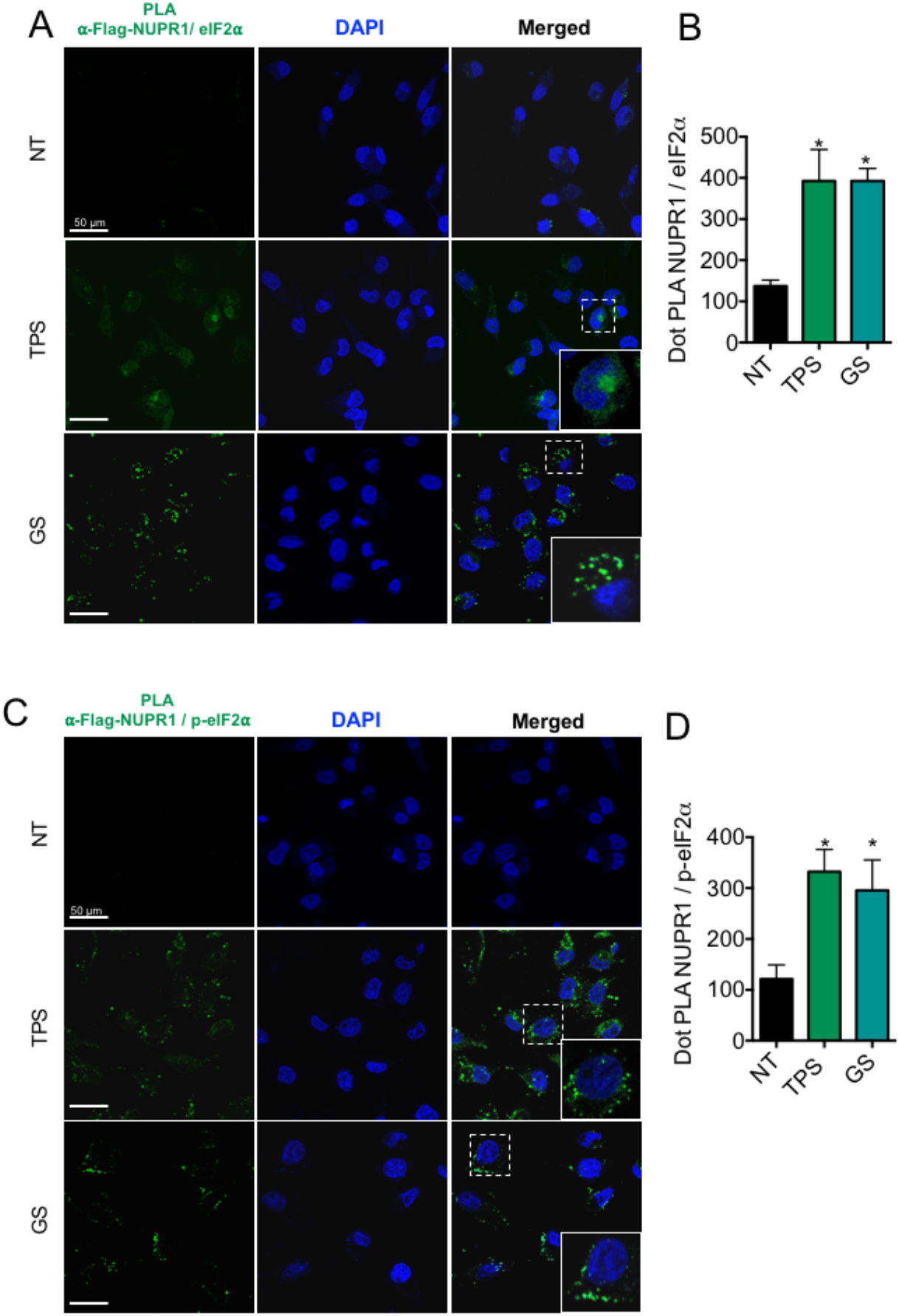
*In situ* proximity ligation assay (PLA) between Flag-NUPR1 and eIF2α/ p-eIF2α confirms the interaction of the proteins. Representative images of PLA experiment for Flag-NUPR1 and eIF2α **(A)** or Flag-NUPR1 and p-eIF2α **(B)** were used to reveal interaction in Control MiaPaCa-2 cells or following either thapsigargin treatment (TPS, 1 µM) or glucose-starved (GS) for 24 hours. **(B-D)** PLA quantification was performed with ImageJ and involved counting the number of green pixels. Data are means of 10 field each containing not less than 150 nuclei. Statistical significance was calculated using ordinary One-way ANOVA values and corrected for multiple comparison’s using Dunnet’s test (\**p ≤* 0.001). Magnification 60*×*.

### NUPR1-depletion enhances eIF2α phosphorylation and delays the translational recovery after stress induction

So far, our data revealed that NUPR1 interacts with eIF2α both in its phosphorylated and unphosphorylated forms, and this interaction may contribute to alleviating cell stress in acinar cells. The maintenance of high levels of p-eIF2α in *Nupr1*^*-/-*^ acini propose a models where NUPR1 would contribute directly or indirectly to the dephosphorylation of eIF2α as a mechanism to allow restoration of protein translation. To test this model, we examined phosphorylation of eIF2α in response to ER stress response in PANC-1 cells (*NUPR1*^*+/+*^) or NUPR1-null cells (*NUPR1*^*-/*-^), generated by CRISPR/Cas9 deletion of *NUPR1* [29]. Treatment with 1 µM of TPS for up to 24 hours showed *NUPR1*^*-/-*^ cells maintained higher levels of p-eIF2α compared to *NUPR1*^*+/+*^ cells (Figure 6A,B). The results prompt us to investigate whether the absence of NUPR1 interferes with expression of downstream effectors of eIF2α signaling, including CHOP and ATF4. Consistent with our *in vivo* pancreatic data, *NUPR1*^*-/*-^ cells showed delayed and reduced expression of ATF4 and CHOP compared to *NUPR1*^*+/+*^ cells, which show increased expression of these markers within 6 hours of TPS treatment. RT-qPCR results confirmed reduced expression of both *CHOP* and *GADD34* mRNAs at all time points in *NUPR1*^*-/*-^ cells (Figure 6D), consistent with a deficit in PERK/eIF2α signaling. At later stages of stress, GADD34 participates in eIF2α dephosphorylation to revert the protein translation shut off. The hindered expression of UPR markers such as CHOP and ATF4 in *in vivo* and *in vitro* NUPR1 loss of function models could be therefore a direct cause of a sustained phosphorylation of eIF2α. By expressing lower levels of GADD34 mRNA expression at all time points, it could be that NUPR1-depleted cells have a possible defect in the eIF2α dephosphorylation process.

**Figure 6.**
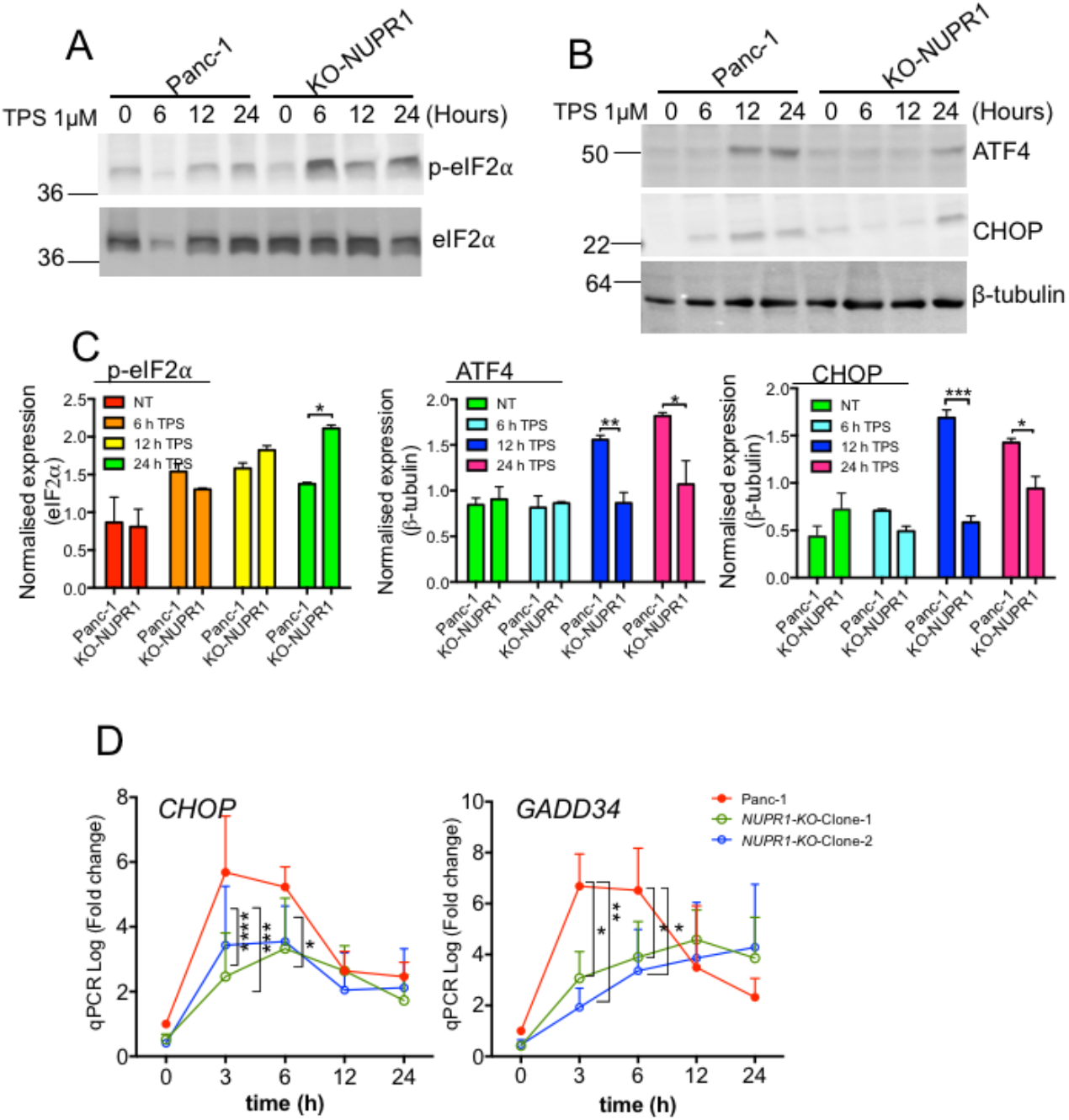
EIF2*α* phosphorylation in *NUPR1*-deficient and matched control PANC-1 cells in response to thapsigargin (TPS) induced ER-stress. **(A)** Analysis of p-eIF2*α* expression following 6, 12 and 24 h TPS treatment in protein extracts of PANC-1 cells with *NUPR1*^*+/+*^ (WT) or following CRISPR-Cas9-mediated deletion of *NUPR1*^*-/-*^ (KO-NUPR1). **(B)** Analysis of ATF4 and CHOP expression following 6, 12 and 24 h TPS treatment in protein extracts of PANC-1 *NUPR1*^*+/+*^ or *NUPR1*^*-/-*^ cells (KO Clone-1). (C) Quantification of A and B using ImageJ software. Mean band intensity plotted ± SEM (n=3). Significant differences were calculated by two-way ANOVA with post hoc Sidak’s test (n=3). **(C)** Analysis by RT-qPCR of *CHOP* and *GADD34* expression in *NUPR1*-deficient and matched control cells after treatment with 1 µM TPS. Significant results between the two genotypes are reported in the graph *(****p* < 0.0001, *\*\*\*p* < 0.001, *\*\*p =* 0.0012, *\*p =* 0.04 at 3, 6, 12 and 24 hours respectively).

Sustained p-eIF2α and low levels of GADD34 should affect translational recovery after stress induction. Therefore, to examine the restoration of protein translation in the absence of NUPR1, we induced the UPR for one hour with TPS (1 µM) in *NUPR1*^*+/+*^ and *NUPR1*^-/-^ PANC-1 clones and assessed the nascent proteins for up to 16 h later using *in situ* click chemistry (Figure 7). To do this, cells were incubated with a puromycin analogue bearing a propargyl group for 1 hour, allowing co-translational incorporation at the C-terminus of nascent polypeptides chains. The incorporated propargyl-puromycin form an *in situ* covalent conjugate by copper catalyzed click reaction with a fluorescently labelled azide (FITC-azide). The reaction enables subsequent analysis of protein synthesis based on FITC fluorescence levels using confocal microscopy and cytometry [30].

**Figure 7.**
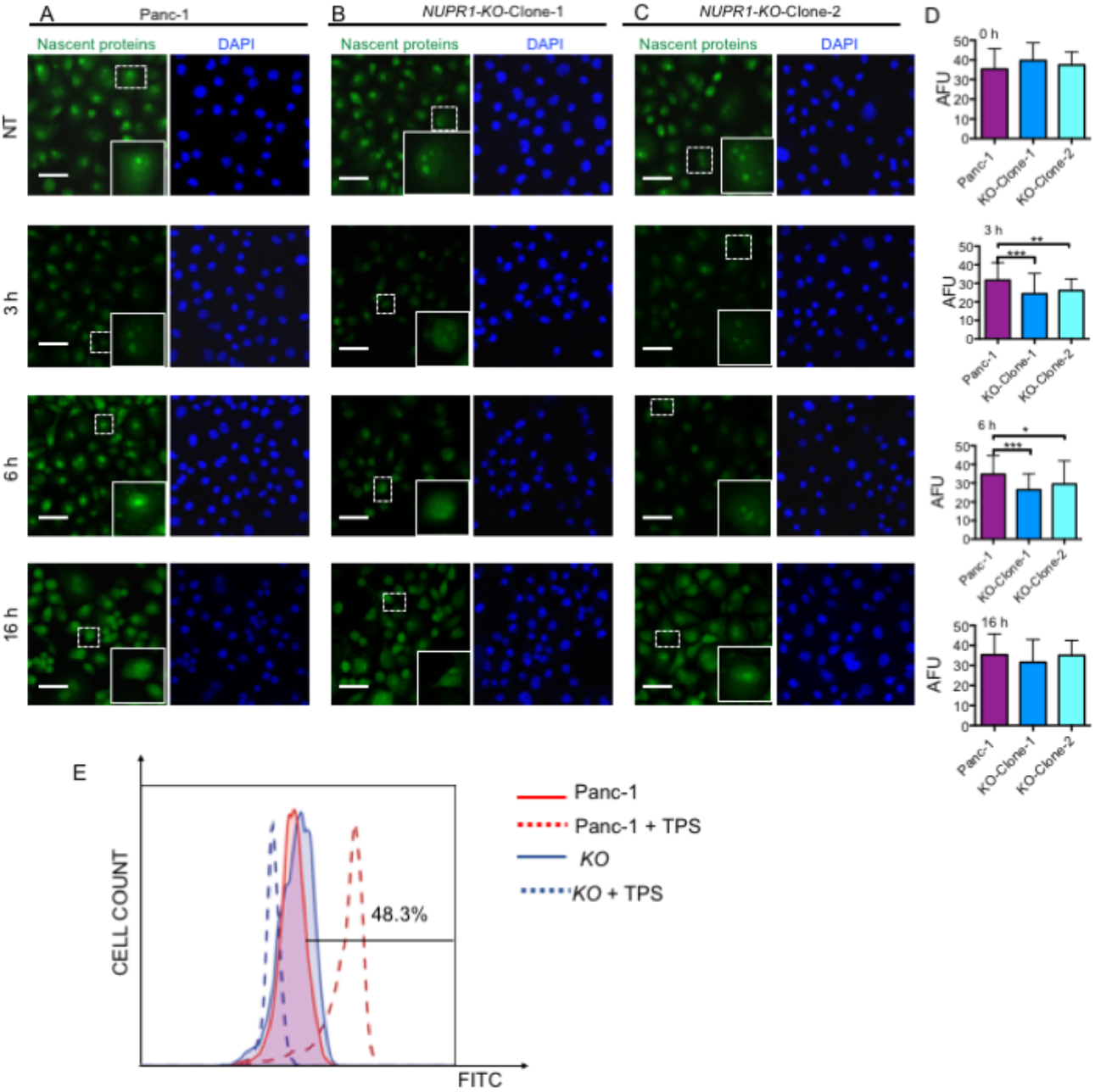
NUPR1 deletion reduces nascent protein production in pancreatic cancer cells after stress induction. **(A-C)** Cultured PANC-1 cells or following CRISPR-Cas9-mediated deletion of *NUPR1* (*NUPR1*^*-/-*^NUPR1-KO-Clone 1 and NUPR1-KO-Clone 2) were treated with vehicle or 1 µM of TPS for one hour. After 3-6 or 16 h, cells were incubated for 1 hour with OP-puromycin. Fluorescence was analysed by confocal microscopy and arbitrary fluorescent intensity (AFU) measured with imageJ **(D)**. Two-way ANOVA with a post hoc Sidak’s test was used to determine statistical significance (\*\**p* = 0.001, \*\*\*\**p*< 0.001). **(E)** Flow cytometry of *de-novo* protein synthesis measured with OP-puromycin. Cells were gated based on forward scatter (FSC) and side scatter (SSC) parameters. The mean fluorescence increases are reported on the X axis for WT or KO PANC-1 cells subjected to click chemistry of integrated OP-puromycin conjugated with FITC-azide. The Y-axis represents the cell count.

NUPR1-depleted cells showed a slower translational recover after stress treatment as shown by a reduction in *de novo* protein synthesis at 3 and 16 hours At three hours after TPS incubation withdrawal, *NUPR1*^*-/-*^ cells displayed lower levels of fluorescence compared to untreated cells (Figure 7A-C and quantification D), suggesting reduced protein translation. Decreased fluorescence was confirmed by two-way ANOVA, which showed an effect of both TPS *(p* < 0.0001) and NUPR1 (*p <* 0.0001), as well as an interaction between the two factors (*p* = 0.001). Six hours post-TPS treatment, *NUPR1*^*+/+*^ PANC-1 cells showed increased fluorescence, which is proportional to increased nascent protein synthesis. In line with sustained eIF2α phosphorylation, both *NUPR1*^*-/-*^ clones maintained lower levels of fluorescence (Figure 7D) until 16 h post pharmacological stress termination. Confocal results were confirmed at 6 hours of TPS treatment by flow cytometry (Figure 7E). Thus, our observations suggest a novel role for NUPR1 in modulating protein synthesis in response to cellular stress.

## Discussion

NUPR1 is rapidly activated in response to a variety of stresses including several ER stress inductors such as serum starvation, cycloheximide, ceramide, staurosporine and CCl_4_ [2,4,6,19,31]. Until now, a direct role for NUPR1’s within the UPR has not been identified. In this study, we identified a completely novel function for NUPR1 in directly interacting with eIF2α/p-eIF2α and affecting the translational machinery regulating ER stress. *NUPR1*-deficient cells showed reduced protein translation following induction of stress both *in vivo* and *in vitro*. This was combined with a lack of ER dilation, which is typically associated with re-activation of general protein translation, suggested NUPR1 is required for restoring protein translation. Using global (mass spectrometry) and targeted (co-IP, PLA) assays, we showed a novel and direct interaction between NUPR1 and eIF2α, and show this interaction occurs in perinuclear regions of the cell. These findings constitute the first evidence of a direct, non-nuclear role for NUPR1 and are consistent with specific deficits observed in the absence of NUPR1, including reduced protein translation and ultrastructural differences in response to induction of the UPR.

The UPR is a critical intracellular signaling pathway maintaining cellular homeostasis by finely tuning responses to various metabolic, oxidative or inflammatory stresses [31]. A maladaptive UPR has been implicated in a variety of metabolic, neurodegenerative, and inflammatory diseases, as well as cancer. The main goal of an acute activation of UPR is to restore the ER homeostasis [12,32] and promote mechanisms directed to reducing misfolded proteins in the ER. These mechanisms include ubiquitination followed by proteasome degradation of misfolded proteins [33,34], autophagy [16,35], and the transitory arrest of protein synthesis and RNA processing that prevents accumulation of misfolded neo-proteins into the ER [16,36]. Efficient activation of these mechanisms allows the cell to survive. Conversely, when these mechanisms are inadequate or sustained, ER-stressed cells initiate programmed cell death [16]. Using several complementary approaches, we have demonstrated NUPR1 affects translation during ER stress, at least in part, by associating with p-eIF2α/eIF2α and potentially promoting dephosphorylation of p-eIF2α, thereby allowing restoration of normal protein synthesis. As a consequence, in NUPR1-deficient cells, protein synthesis is almost completely arrested correlating to prolonged phosphorylation of eIF2α. Decreased eIF2α activity, would result in a reduced ER stress response. While we have not shown that NUPR1 interaction affects phosphorylation status of eIF2α, the absence of NUPR1 leads to maintained p-eIF2α and reduced protein translation, consistent with such a function.

It is likely that NUPR1’s interaction with -eIF2α is not the only role it plays in the UPR and protein translation. The range of NUPR1 interacting partners identified by MS analysis suggests its potential in regulating numerous functions during cell stress, including several transcriptional regulators, one of such functions could be indeed the transcriptional regulation. We chose to focus on eIF2*α* as it is a critical translational regulator that mediates the ISR. Upon ER-stress induction, the first cellular event in the UPR cascade involves eIF2α phosphorylation that prompts a protein synthesis shut off to re-establish the pre-stress proteostasis. Both the phosphorylation of eIF2α and the subsequent recovery of protein translation are important stages in a complete UPR. In NUPR1 deficient cells we observed an elevated and prolonged phosphorylation of eIF2α associated with a reduced and delayed induction of UPR downstream effectors CHOP, ATF4 and GADD34. The reduced expression and function of ATF4 (a direct regulator of *Chop* and *Gadd34* transcription) is somewhat surprising as *Atf4* generally escapes the translation block normally bestowed by p-eIF2α. However, it is likely that NUPR1 deficiency affects multiple factors within the UPR, possibly through transcriptional mechanisms. Indeed, we observed deficits in IRE1 signaling suggesting a more widespread effect of NUPR1 deficiency on the UPR. Another possible mechanism involves GADD34. In the later stages of the ER-stress response GADD34 promotes the dephosphorylation of eIF2α thereby creating a negative feedback loop to release the transient protein synthesis block. Loss of GADD34 has been previously associated with a prolonged interruption in protein synthesis [32] and decreased expression of UPR downstream regulators. Since we show specific decreases in GADD34 expression both *in vitro* and *in vivo*, it is possible that this decrease contributes to maintained p-eIF2α levels as well. However, our MS data did not demonstrate a direct interaction of NUPR1 with GADD34. Noteworthy, in light of our findings it could also be speculated that the prolonged phosphorylation of eIF2α in absence of NUPR1 is due to an improper eIF2α dephosphorylation. The proposed observation fits with the model that the eIF2α phosphorylation leads to a reduced expression of CHOP in absence of NUPR1. Moreover, we identify a direct interaction between NUPR1 and eIF2α supported by our MS, biochemical and PLA indicate that NUPR1 in addition to transcriptional regulation of CHOP and GADD34, could potentially contribute to the regulation of eIF2α phosphorylation through direct interaction with proteins involved in eIF2α phosphorylation or dephosphorylation. However, the exact mechanism clearly warrants further investigation. While our data demonstrates NUPR1 functions influence protein translation through interacting with factors involved in translation initiation and regulation, it is likely that NUPR1 affects the UPR through multiple mechanisms.

In conclusion our findings demonstrate a novel role for NUPR1 during ER stress where it participates in the regulation of the UPR and more broadly the integrated stress response by participating in the transcriptional and translational regulation by interacting with eIF2α (Figure 8). Thus, collectively, our data supports an essential role of NUPR1 during ER stress.

**Figure 8.**
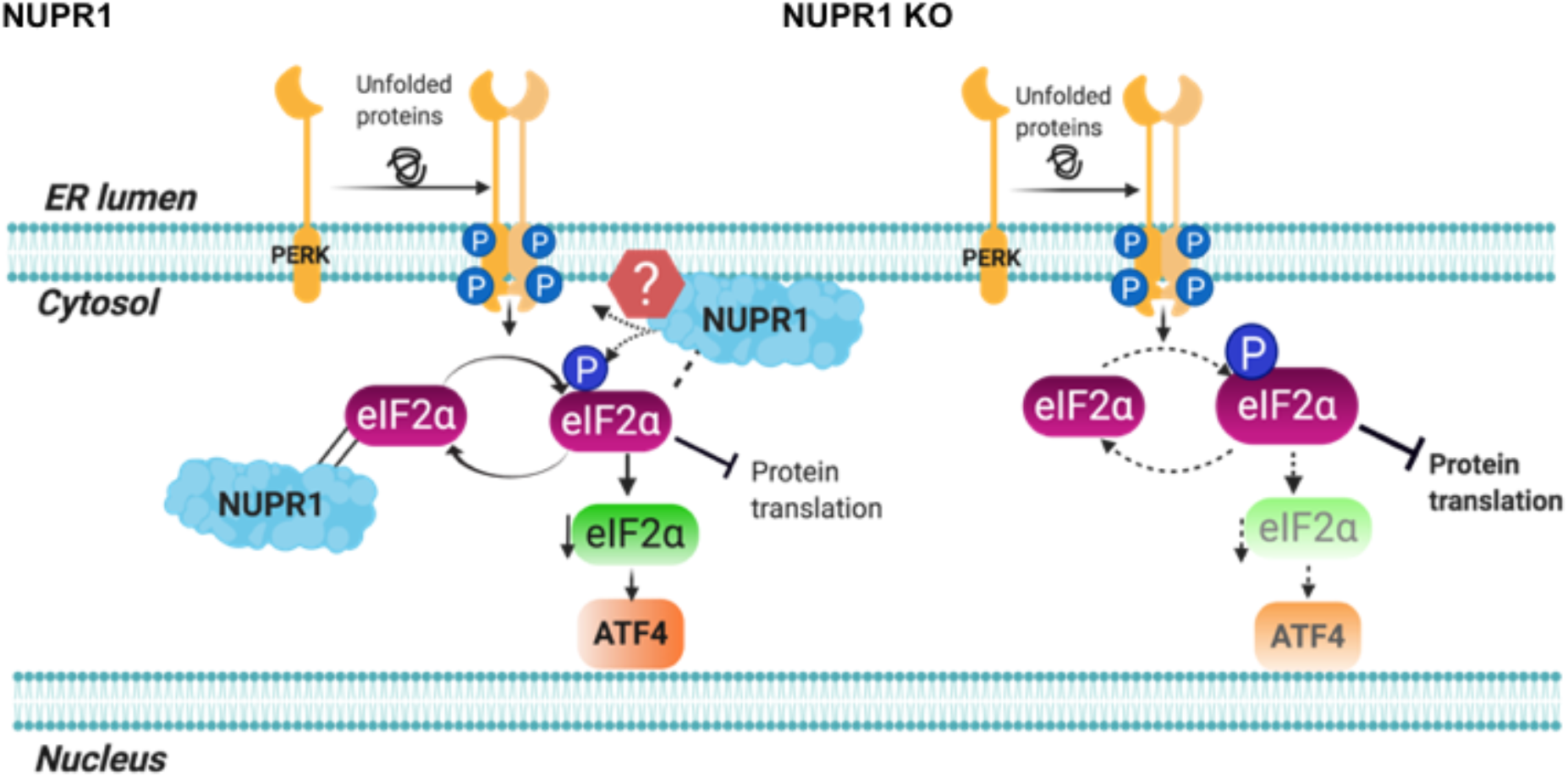
The stress-induced protein NUPR1 orchestrates protein translation during ER-stress by interacting with eIF2*α*: NUPR1 by interacting with the eIF2*α* participates in the translational regulation after stress. In the absence of NUPR1 the phosphorylation of eIF2*α* is maintained for longer and all the expression of downstream regulators of ER is impaired.

## Materials and methods

### Study approval

All experimental protocols were carried out in accordance with the nationally approved guidelines for the treatment of laboratory animals. All experimental procedures on animals were approved by the Comité d’éthique de Marseille numéro 14 (C2EA-14) in accordance with the European Union regulations for animal experiments.

### Mouse strains and Tissue collection

For all the *in vivo* experiments we used *Nupr1*^*-/-*^ mice bear a homozygous deletion of exon 2 [33]. Mice were used between the 5 and 16 weeks of age and as control we used their mating littermates. Animals were kept in the Experimental Animal House of the Centre de Cancérologie de Marseille (CRCM) of Luminy. After sacrifice by cervical dislocation, pieces of pancreas were collected and frozen in cold isopentane for further analysis or directly homogenized in 4 M guanidium isothiocyanate lysis buffer for efficient pancreatic RNA extraction according with Chirgwin et al’s procedure [34].

### Tunicamycin injections

Mouse of 5-8 weeks of age were injected intraperitoneally (IP) with 1 µg/g of Tunicamycin (Sigma) using 150 mM of D-Glucose as vehicle. After the reported time point (up to 36 h) mice were sacrificed and organs collected.

### Cell culture

MiaPaCa-2 and PANC-1 cells were purchased from ATCC and cultures in DMEM medium (Dulbecco’s modified Eagle’s medium, Gibco, Life Technologies, Carlsbad, CA, USA), supplemented with 10% fetal bovine serum (Lonza, Basel, Switzerland) and cultured at 37°C and 5% CO_2_. Cells were used in all experiments below 20 passages.

### Preparation or protein lysates and Western Blotting

Forty mg of frozen tissue were cut on dry ice and homogenized with Precellys^®^ (Bertin instruments) in 300 µL of ice-cold RIPA buffer (10 mM Tris-Cl (pH 8.0), 1 mM EDTA, 0.5 mM EDTA, 1% Triton X-100, 0.1% sodium deoxycholate, 0.1% SDS, 140 mM NaCl), complemented with 0.5 µg/g of protease inhibitor kit (Sigma), 200 µM of Na_3_VO_4_, 1 mM of PMSF and 40 mM of β-glycerophosphate. After homogenization, the supernatant was cleared by centrifugation for 30 min at 14.000 rpm at 4°C. Protein lysate content was quantified by micro BCA assay (Thermo-Fisher Scientific) and 40 µg of protein were resolved by SDS-PAGE, transferred to nitrocellulose membrane for 1 or 2 hours. Membranes were next blocked with TBS (tris-buffered saline) in 5% BSA and blotted overnight with primary antibody (1:500 dilution in TBS 5% BSA). After washes, the membrane was incubated with HPR-conjugated secondary antibody (Boster, Pleasanton CA, USA) for 1 hour at room temperature (1:5000 dilution in TBS-5% BSA). Subsequently, the membrane was washed and revealed with ECL (enhanced chemo-luminescence). Chemiluminescent signal was detected in a G-Box (Syngene). The following antibodies were used: rabbit monoclonal ATF4 (D4B8), mouse monoclonal CHOP (L63F7), rabbit monoclonal XBP1s (E9V3E), rabbit monoclonal, BiP (C50B12), rabbit monoclonal eIF2α XP^®^ (D7D3), rabbit monoclonal Phospho-eIF2α XP^®^ (Ser51) (D9G8), and rabbit monoclonal Phospho-PERK (Thr980) were from Cell Signaling Technology;; (G.305.4), rabbit polyclonal antibody Phospho-IRE1 alpha (Ser724) (PA1-16927) is from Thermo Fisher;; and mouse monoclonal β-actin (#A5316) is from Sigma. Quantification of signal was performed using ImageJ software. Mean band intensity plotted over the intensity of eIF2*α* ± SEM (n=3) and unpaired Student’s *t*-test was used for statistical analysis.

### Preparation of RNA and RT-qPCR

RNA from murine pancreata was extracted after 16 h of intraperitoneal injections of Tunicamycin following the Chirgwin procedure [34]. Total RNA from cells was obtained using RNAeasy kit (Quiagen) following manufacturer’s instruction. The RNA integrity was assessed with Agilent 2100 Bioanalyzer with the RNA 6000 Nano chip Kit. The RNAs were reverse-transcribed using GO Script kit (Promega) following manufacturer’s instruction. RT-qPCR were performed using Aria mix using Promega reagents. Primer sequences are described in Supplementary Table 4. mRNAs were quantified relative to *Rpl0*. Data are presented in the graphs as Log Fold Change compared to *Nupr1*^*+/+*^ controls (mice IP injected with vehicle, 150 mM D-Glucose) levels of expression. Significant differences were calculated using two-way ANOVA with *post hoc* Sidak test (n=6).

### Transmission Electron Microscopy

Mice were perfused with 4% cold PFA and 2.5% glutaraldehyde. Pancreatic tissue was then immersed overnight in 0.1M Soresen buffer, post-fixed in 1% osmium tetroxide, and in bloc stained with 3% uranyl acetate. The tissue was dehydrated with increasing concentrations of ethanol on ice and acetone before being embedded in Epon. Ultrathin sections (70 nm) were prepared using a Leica UCT Ultramicrotome (Leica, Austria) and stained with uranyl acetate and lead citrate and deposited on formvar-coated slot grids. The grids were observed in an FEI Tecnai G2 at 200 KeV and acquisition was performed on a Veleta camera (Olympus, Japan).

### NUPR1 expression vector transfection

MiaPaCa-2 cells were seeded in 12-well plates and transfected with 3 µg of DNA (NUPR1-GFP, NUPR1-Flag or control vector) and 3 μL of Lipofectamine 3000 Transfection Reagent (Thermo Fisher Scientific) per well. Cells were assayed after 24 h post-transfection.

### Co-Immunoprecipitation

MiaPaCa-2 cells, expressing GFP-NUPR1, were plated in 10 cm^2^ dishes. When reached 80% confluence were treated with either 1 µM of Thapsigargin or glucose starved for 24 h. After that time cells were lysed on ice by using HEPES based lysis buffer containing proteases inhibitor cocktail (1:200) (Sigma P8340). Lysates were cleared for 10 min at 14000 rpm at 4°C and protein concentration of the supernatant was determined by using Protein Assay (BioRad). The co-immunoprecipitation was performed using GFP trap^®^ beads (Chromotek) following manufacturer’s protocol or rabbit monoclonal antibody specific for eIF2α XP^®^ (D7D3). Immunoprecipitates were pelleted, washed with lysis buffer three times, and then PBS. The resultant proteins were denatured and blotted against eIF2α, p-eIF2*α* and anti GFP. To perform the reverse co-IP, MiaPaCa-2 cells expressing Flag-NUPR1 or Flag-GFP were grown to 70% confluence and treated as described above. Cells were lysed on ice using HEPES based lysis buffer containing 10 mM NEM (N-Ethylmaleimide, Sigma 04259) and a proteases inhibitor cocktail (1:200;; Sigma P8340). Lysates were centrifuged for 10 min at 14, 000 rpm at 4°C. Protein concentration of the supernatant was determined using Protein Assay (Bio-Rad), and equal amounts of protein incubated with 30 µl of anti-Flag M2 coated beads under rotation for 2 h at 4°C. Beads were then washed three times with cold lysis buffer and proteins were eluted using 250 µl ammonium hydrogen carbonate buffer containing 0.1 µg/µl of Flag peptide for 90 min at 4°C while rotating. After a short spin, the supernatant was recovered by using a Hamilton syringe. Eluted proteins were collected and analyzed by mass spectrometry.

### Click chemistry and fluorescence detection

PANC-1 and modified PANC-1 cells were grown on glass coverslips in DMEM supplemented with 10% bovine calf serum. Cell were treated with 1 µM Thapsigargin (TPS) for 1 h then incubated in fresh media without TPS. After 3, 6 or 16 hours, O-propargyl puromycin (OP-puro) was added to cells in complete culture medium for 1h. Cells were then washed with PBS and fixed with 4% PFA in PBS. After fixation cells were permeabilized with 0.3% Triton-X in PBS. Following removal of detergent by PBS washes, CuAAC detection of OP-puro incorporated into nascent protein was performed by reacting the fixed cells for 1 h at room temperature with 20 μM FITC azide, as previously described [35]. After the click chemistry reaction, coverslips were washed several times with TBST, counterstained with Hoechst, and mounted in standard mounting media. The stained cells were imaged by LSM 510 META confocal microscope (Zeiss) and on a Nikon Eclipse 90i fluorescence microscope. Stained cells were also quantified by Flow-cytometry in a MACSQuant-VYB (Miltenyi Biotec, Surrey, UK). Data analysis was carried out by using the FlowJo software. The intensity of the fluorescent OP-puro stain in single cells was quantified by imageJ. Statistical significance was calculated by using two-way ANOVA and corrected for Sidak s test.

### Proximity ligation assay

MiaPaCa-2 transfected with Flag-NUPR1 were treated with TPS (1µM), or glucose starved. After the indicated time points cells were washed twice in PBS, fixed, washed twice again, permeabilized in PBS/0.1% Triton X-100, and saturated with blocking solution for 30 min before immune-staining with Duolink by using PLA Technology (Sigma-Aldrich) following the manufacturer’s protocol. Slides were processed for *in situ* PLA by using sequentially the Duolink *in situ* Detection Reagents Green, Duolink In Situ PLA Probe Anti-Mouse MINUS, and Duolink In Situ PLA Probe Anti-Rabbit PLUS. The following antibodies were used: rabbit monoclonal eIF2α-XP^®^ (D7D3, from Cell Signaling Technology), rabbit monoclonal Phospho-eIF2α XP^®^ (Ser51) (D9G8, from Cell Signaling Technology), mouse monoclonal antibody anti-FLAG M2 (from Sigma-Aldrich). In these experiments, green fluorescence corresponds to the PLA-positive signal, and it indicates that the two molecules belong to the same protein complex. Blue fluorescence corresponds to nuclei (so-called DAPI staining). Protein overexpression was used to obtain a clearer and better signal. Preparations were mounted using Prolong Gold antifade reagent (Invitrogen) and image acquisition was carried out on an LSM 510 META confocal microscope (Zeiss) and on a Nikon Eclipse 90i fluorescence microscope.

### Mass Spectrometry Analysis

Protein extracts were loaded on NuPAGE 4–12% Bis-Tris acrylamide gels according to the manufacturer’s instructions (Invitrogen). Running was stopped as soon as proteins stacked in a single band. Protein-containing bands were stained with Imperial Blue (Pierce), cut from the gel, and digested with high-sequencing-grade trypsin (Promega, Madison, WI) before mass spectrometry analysis according to Shevchenko et al. (44). Mass spectrometry analysis was carried out by LC–MS/MS using an LTQ-Velos-Orbitrap or a Q Exactive Plus Hybrid Quadrupole-Orbitrap (Thermo Electron, Bremen, Germany) coupled online with a nanoLC Ultimate3000RSLC chromatography system (Dionex, Sunnyvale, CA). Five microliters corresponding to 1/5 of the whole sample were injected in triplicate on the system. After sample preconcentration and washing on a Dionex Acclaim PepMap 100 C18 column (2 cm x 100 μm i.d. 100 Å, 5 μm particle size), peptides were separated on a Dionex Acclaim PepMap RSLC C18 column (15 cm x 75 μm i.d., 100 Å, 2 μm particle size) at a flow rate of 300 nL/min, a two-step linear gradient (4–20% acetonitrile/H2O;; 0.1% formic acid for 90 min and 20–45% acetonitrile/H2O;; 0.1% formic acid for 30 min). For peptides ionization in the nanospray source, voltage was set at 1.9 kV and the capillary temperature at 275 °C. All samples were measured in a data-dependent acquisition mode. Each experiment was preceded by a blank run to monitor system background. The peptide masses were measured in the LTQ-velos-orbitrap in a survey full scan (scan range 300–1700 m/z, with 30 K FWHM resolution at m/z = 400, target AGC value of 1.00 × 106, and maximum injection time of 200 ms). In parallel to the high-resolution full scan in the Orbitrap, the data dependent CID scans of the 10 most intense precursor ions were fragmented and measured in the linear ion trap (normalized collision energy of 35%, activation time of 10 ms, target AGC value of 1 × 104, maximum injection time 100 ms, and isolation window 2 Da). Parent masses obtained in Orbitrap analyzer were automatically calibrated on 445.1200 locked mass. Dynamic exclusion was implemented with a repeat count of 1 and exclusion time of 30 s.

In the Q Exactive Plus Hybrid Quadrupole-Orbitrap, the peptide masses were measured in a survey full scan (scan range 375-1500 m/z, with 70 K FWHM resolution at m/z=400, target AGC value of 3.00 × 106 and maximum injection time of 100 ms). Following the high-resolution full scan in the Orbitrap, the 10 most intense data-dependent precursor ions were successively fragmented in higher energy collisional dissociation (HCD) cell and measured in Orbitrap (normalized collision energy of 25 %, activation time of 10 ms, target AGC value of 1.00 × 103, intensity threshold 1.00 × 104 maximum injection time 100 ms, isolation window 2 m/z, 17.5 K FWHM resolution, scan range 200 to 2000 m/z). Dynamic exclusion was implemented with a repeat count of 1 and exclusion time of 20 s.

### Mass Spectrometry Data Analysis

Raw files generated from mass spectrometry analysis were processed using Proteome Discoverer 1.4.1.14 (Thermo Fisher Scientific). This software was used to search data via in-house Mascot server (version 2.3.0;; Matrix Science, London, U.K.) against the Human database subset of the SwissProt database (version 2017.03, 20184 human entries). A database search was done by using the following settings: a maximum of two trypsin miscleavage allowed, methionine oxidation and protein N-acetylation as dynamic modifications, and cysteine carbamido-methylation as fixed modification. A peptide mass tolerance of 6 ppm and a fragment mass tolerance of 0.8 Da were allowed for search analysis. Only peptide identified with a FDR < 1% were used for protein identification.

To calculate the confident score (from 0 to 100%) for NUPR1-interacting proteins identified by MS, we used a formula derived from Bonacci et al. [36] based on peptide number count: K=total peptide number in control IP;; V=total peptide number in NUPR1 IP;; Conf = ((2V)^2^/(1+(2V)+(2K)^2^)*100 - 100/(1+(2(V-K));; = 0 if <0;; Values above 50 are usually considered to be confident.

## Acknowledgements

The work was supported by La ligue Contre le Cancer, INCa, Canceropole PACA and INSERM. The electron microscopy experiments were performed in the PiCSL-FBI core facility (IBDM, AMU-Marseille).

## Figure Legends

Figures were realized with Biorender

## Notes

### Competing Interest Statement

The authors have declared no competing interest.

